# Selective pressures on human cancer genes along the evolution of mammals

**DOI:** 10.1101/388421

**Authors:** Alberto Vicens, David Posada

**Affiliations:** Department of Biochemistry, Genetics and Immunology, University of Vigo, 6310, Vigo, Spain; Biomedical Research Center (CINBIO), University of Vigo, 36310 Vigo, Spain; Galicia Sur Health Research Institute, 36310 Vigo, Spain

**Keywords:** positive selection, somatic evolution, germline evolution, *dN/dS*

## Abstract

Cancer is a disease of the genome caused by somatic mutation and subsequent clonal selection. Several genes associated to cancer in humans, hereafter cancer genes, also show evidence of (germline) positive selection among species. Taking advantage of a large collection of mammalian genomes, we systematically looked for statistically significant signatures of positive selection using *dN/dS* models in a list of 430 cancer genes. Among these, we identified 63 genes under putative positive selection in mammals, which are significantly enriched in processes like crosslinking DNA repair. We also found evidence of a higher incidence of positive selection in cancer genes bearing germline mutations, like BRCA2, where positively selected residues are physically linked with known pathogenic variants, suggesting a potential association between germline positive selection and risk of hereditary cancer. Overall, our results suggest that genes associated with hereditary cancer have less selective constraints than genes related to sporadic cancer. Also, that the adaptive evolution of human cancer genes in mammals has been most likely driven by adaptive changes in important traits not directly related to cancer.

## 1. Introduction

Cancer is a genomic disease caused by mutations in genes that control normal cell functions, in particular growth and division. A fundamental goal of cancer genomics is to identify mutations that confer a selective advantage to the cell and increase survival and proliferation, so-called driver mutations, as well as the genes carrying the driver mutations in each tumor, known as driver genes or cancer genes. Although traditionally the focus has been put on somatic driver mutations, those that appear during an individual lifetime as cells divide and grow, there are also germline mutations in the human population that predispose to cancer [1]. Today, more than 500 cancer genes have been identified, of which approximately 90% contain somatic mutations and 20% bear germline mutations [2–4].

Several studies have identified a number of human cancer genes undergoing positive selection across species [5]. Clark et al. (2003) found strong evidence of positive selection on oncogenes and tumor suppressor genes in the chimpanzee lineage [6]. Nielsen et al. (2005) identified an elevated number of tumor suppressor and apoptosis genes under strong positive selection in humans and chimpanzees [7]. Subsequent genome-wide screenings in mammals unveiled positive selection in genes with roles in immunity and reproduction, but also related with apoptosis and cancer [8,9]. Given the recurrent observation of positive selection acting on human cancer genes across species, these authors have proposed that evolutionary forces affecting organismal fitness, such as sexual selection, pathogen-host interactions or maternal-foetal conflict, could also lead to increased cancer risk in humans as a pleiotropic effect [5,7,9,10]. If this hypothesis is correct, genes under positive selection during the evolution of species should have somehow a higher contribution to cancer risk than those genes subjected to strong purifying selection. Another intriguing question is whether selective constraints in mammals differ across genes involved in different human tumor types. For example, since blood and bone marrow cancers, such as leukemia and lymphoma, are promoted by alterations in the immune system, we would expect stronger positive selection on genes associated with these cancers due to the faster adaptation of immunity-related genes [7,9].

Here, we carried out a comprehensive analysis of the evolution of 430 human cancer genes in mammals to address these questions in more detail. Using the ratio of number of non-synonymous substitutions per site to the number of synonymous substitutions per site (*dN/dS)* [11], we identified 63 genes under putative positive selection, significantly enriched in double-strand DNA break repair, and associated with hereditary cancer and recessive syndromes.

## 2. Materials and Methods

### 2.1 Cancer genes

We retrieved a collection of single-copy 574 genes associated with human cancer from the Cancer Gene Census (CGC) project of the COSMIC repository [3] (accessed on March 5th 2018). We only collected genes classified into Tier 1, which refers to genes with a documented activity relevant to cancer. The catalogue of the retrieved cancer genes, along with information about their function and associated mutations, can be accessed in Table S1.

### 2.2 Sequence data collection

We downloaded a representative sequence for each cancer gene from the Ensembl Genes database (Release 91, human genome version GRC) using BioMart [12] (accessed on March 6th 2018). For each human gene, we chose a single isoform based on the following criteria, in order: GENCODE validation, APRIS annotation as principal 1, best transcript support level (TSL), and longer transcript. We discarded 22 genes whose best TSL was less than one (Table S2). Using the selected human isoform as reference, we downloaded the corresponding orthologues from 32 mammalian genomes (Table S3, Figure S1) using the Bioconductor package BiomaRt [13]. When more than one ortholog was obtained for a given species, we chose the one with the best orthology confidence score. We discarded 17 genes for which less than 15 mammal orthologues were found (Table S2), so the final number of retrieved ortholog groups was 535 (Table S4).

### 2.3 Multiple sequence alignment

We aligned the coding sequences for each ortholog group using MACSE [14], a program that accounts for frameshifts and stop codons. The resulting multiple sequence alignments were further refined with TrimAl [15], removing taxa and sites with more than 60% gaps across rows and columns, respectively. After trimming, we discarded 71 genes that contained less than 10 orthologues (Table S2) in order to maximize the statistical power for the selection analyses, ending up with a list of 464 genes.

### 2.4 Estimation of phylogenetic trees

We inferred maximum likelihood trees for the 464 genes using RAxML-NG [16]. All reconstructions were performed using the general time reversible substitution model [17] with gamma-distributed rate variation among sites [18]. For each gene, we obtained 10 starting trees using randomized stepwise addition parsimony. We assessed nodal support using 100 bootstrap replicates [19]. To minimize the impact of estimation errors and/or incomplete lineage sorting in subsequent analyses, we discarded 27 genes whose estimated tree topologies were quite distinct (normalized Robinson-Foulds (RF) distance >= 0.6) from a species tree assembled for 19 mammals with a well-known phylogenetic position (Figure S1). We calculated the RF distances with ETE3-compare [20].

### 2.5 Codon-based selection models

We estimated nonsynonymous (*dN*) and synonymous (*dS*) substitution rates using the program codeml of the PAML package v4.9c [21] along 437 mammal gene trees. Because dS saturation decreases the power for detecting positive selection in codon-based models [22], we further discarded seven genes with an estimated dS > 15 (Table S2). To estimate the global *dN/dS* ratios for each of the remaining 430 genes we used the one-ratio (M0) model, which assumes the same *dN/dS* for all branches in the gene tree and across sites. To identify genes under putative positive selection, we compared different site-models using likelihood ratio tests (LRTs): M1a (neutral) vs. M2a (selection) and M8 (beta selection) vs. M8a (beta neutral) [23,24]. The resulting p-values were adjusted for multiple testing using the Benjamini-Hochberg procedure [25] with a family-wise significance level of 0.01. For those genes in which the LRT was significant (i.e., positively selected cancer genes), we considered as positively selected sites (PSSs) those with a Empirical Bayes (BEB) posterior probability > 0.95 of having a dN/dS > 1 under the M2a or M8 models [26].

### 2.6 Gene ontology enrichment analysis

To identify enriched Gene Ontology (GO) terms in the genes under putative positive selection, we used GOrilla [27]. We compared the list of 63 positively selected genes with a background list of the initial 574 cancer genes. We searched for GO terms in the three available ontologies: biological process, cellular component and molecular function.

### 2.7 Pathogenic yermline mutations

We retrieved the list of pathogenic germline variants for the BRCA2 gene from the study published by the TCGA PanCanAtlas Germline Working Group [28].

### 2.8 Comparison ofdN/dS ratios across COSMIC categories

We compared the *dN/dS* ratios obtained across five different CGC-COSMIC classifications: mutation type, inheritance, tissue type, cancer role and chromosome type. To test for significant dN/dS differences between and among categories, we performed ANOVAs (for multiple comparisons) and t-tests (for pairwise comparisons) using the ggpubr package [29] for R [30]. We adjusted the *p-value* for multiple pairwise comparisons as described above. To compare the proportion of genes under putative positive selection across groups, we applied the chi-squared test (*p* < 0.01) function (chisq.test) implemented in R.

## 3. Results

After multiple processing steps and stringent criteria (see Materials and Methods), we finally assessed the selective pressures along the mammal phylogeny on 430 human cancer genes. Multiple sequence alignments included 11-32 taxa and were 108-4984 nt long (Table S5).

### 3.1 Long-term selective pressures on human cancer genes

The mean *dN/dS* for all cancer genes examined was 0.12 (Figure 1), consistent with the idea that germline evolution in mammals is strongly dominated by purifying selection *(dN/dS* << 1) [31] and particularly in cancer genes [32]. The LRTs among site-specific *dN/dS* models detected 63 genes under (putative) positive selection (14.65% of the 430 tested genes) after correcting for multiple testing *(p-adj* < 0.01) (Table 1, S5). Of these, 32 were significant for both M1a vs. M2a and M8a vs. M8 comparisons. All 63 genes showed at least one positively selected site (PSS) (Table 1). Four genes yielded more than 20 PSSs under the M8 model: PTPRC (52 PSSs), BRCA2 (25 PSSs), NIN (25 PSSs) and COL1A1 (21 PSSs) (Table 1). All genes with a global *dN/dS* > 0.4 resulted in significant LRTs (Figure 1).

**Figure 1.**
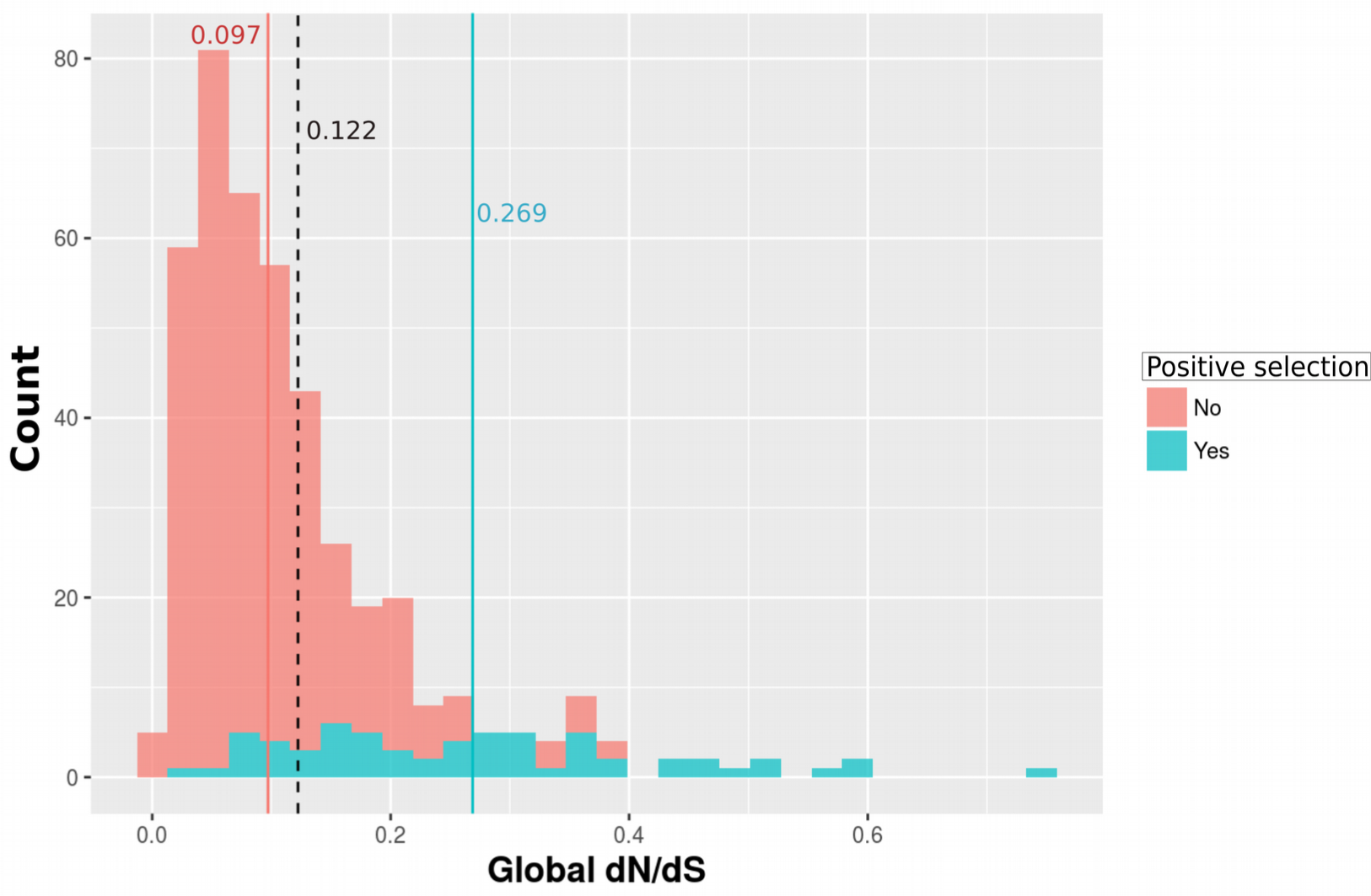
Distribution of global *dN/dS* estimates. Dashed, red and blue vertical lines indicate mean *dN/dS* for all, positively selected, and not positively selected genes, respectively.

**Table 1.** List of cancer genes showing evidence of positive selection.

| Gene | Function | *dN/d*<br>S | *p-adj*<br>M2a-M1a | *p-adj*<br>M8-M8a | PSS<br>M2a/M8 |
| --- | --- | --- | --- | --- | --- |
| ARHGEF12 | regulation of RhoA GTPase | 0.178 | 0.033 | 0.006 | 0/9 |
| B2M | major histocompatibility complex antigen | 0.260 | 0.009 | 0.112 | 1/1 |
| BLM | basic helix-loop-helix transcription factor;nucleic acid binding | 0.264 | 0.000 | 0.000 | 3/2 |
| BRAF | protein kinase; transduction of mitogenic signals | 0.079 | 0.001 | 0.000 | 1/5 |
| BRCA2 | damaged DNA-binding protein | 0.468 | 0.000 | 0.000 | 16/25 |
| BRIP1 | DNA helicase | 0.286 | 0.000 | 0.000 | 4/0 |
| BUB1B | Mitotic checkpoint serine/threonine-protein kinase | 0.267 | 1.000 | 0.000 | 1/2 |
| CASP8 | cysteine protease;protease inhibitor | 0.363 | 0.000 | 0.000 | 10/9 |
| CD274 | immunoglobulin receptor superfamily;membrane-bound signaling molecule | 0.465 | 0.000 | 0.000 | 6/6 |
| CD79A | immunoglobulin receptor superfamily | 0.253 | 0.000 | 0.000 | 5/7 |
| CD79B | immunoglobulin receptor superfamily | 0.275 | 0.000 | 0.005 | 2/3 |
| CDH1 | Cadherin. Cell-cell adhesion | 0.193 | 0.000 | 0.151 | 4/5 |
| CHEK2 | Serine/threonine-protein kinase | 0.175 | 0.000 | 0.023 | 2/2 |
| COL1A1 | Collagen component | 0.276 | 0.000 | 0.000 | 16/21 |
| COL2A1 | Collagen component | 0.132 | 0.000 | 0.035 | 2/11 |
| CSF3R | cytokine;defense/immunity protein | 0.278 | 1.000 | 0.000 | 3/3 |
| CXCR4 | host-virus interaction | 0.061 | 0.000 | 0.054 | 2/4 |
| DDB2 | damaged DNA-binding protein | 0.298 | 1.000 | 0.001 | 1/2 |
| DDIT3 | Cell cycle arrest and apoptosis | 0.190 | 1.000 | 0.009 | 1/1 |
| DICER1 | endodeoxyribonuclease | 0.087 | 0.000 | 1.000 | 3/9 |
| EGFR | growth factor receptor | 0.094 | 0.000 | 0.382 | 3/5 |
| EML4 | microtubule dynamics | 0.144 | 0.001 | 0.342 | 0/4 |
| EPS15 | G-protein modulator;calcium-binding protein;membrane traffic protein | 0.207 | 1.000 | 0.000 | 5/9 |
| ERCC5 | DNA repair | 0.321 | 0.000 | 0.000 | 3/1 |
| FANCA | Fanconi Anemia group; DNA repair | 0.361 | 1.000 | 0.000 | 0/7 |
| FANCC | Fanconi Anemia group; DNA repair | 0.445 | 0.000 | 0.000 | 1/11 |
| FANCD2 | Fanconi Anemia group; DNA repair | 0.348 | 0.000 | 0.000 | 7/8 |
| FANCE | Fanconi Anemia group; DNA repair | 0.393 | 0.000 | 0.000 | 4/9 |
| FANCG | Fanconi Anemia group; DNA repair | 0.500 | 0.000 | 0.000 | 4/7 |
| FAS | acyltransferase;dehydrogenase;esterase;ligase;methyltransferase | 0.601 | 0.000 | 0.000 | 15/15 |
| FCRL4 | B-cell receptor signalling | 0.601 | 0.000 | 0.000 | 8/9 |
| FLT3 | Receptor-type tyrosine-protein kinase | 0.144 | 0.000 | 0.000 | 5/5 |
| IL2 | T-cell proliferation and regulation of the immune response. | 0.748 | 0.000 | 0.000 | 2/3 |
| IL7R | type I cytokine receptor | 0.446 | 0.046 | 0.008 | 0/5 |
| KDM6A | transcription factor | 0.120 | 0.043 | 0.006 | 0/3 |
| KTN1 | Kinesin-driven vesicle motility | 0.211 | 0.000 | 0.000 | 5/6 |
| LYL1 | transcription factor;nucleic acid binding | 0.151 | 1.000 | 0.008 | 1/1 |
| MED12 | nucleic acid binding;transcription cofactor | 0.097 | 0.000 | 0.000 | 1/15 |
| MLLT6 | nucleic acid binding;zinc finger transcription factor | 0.130 | 0.032 | 0.002 | 0/4 |
| NCOA4 | androgen receptor signalling | 0.312 | 0.000 | 0.000 | 4/4 |
| NFE2L2 | transcription factor; response to oxidative stress | 0.236 | 1.000 | 0.002 | 0/3 |
| NIN | centrosome localization | 0.307 | 0.000 | 0.000 | 11/25 |
| NTRK1 | Tyrosin-kinase; neurogenesis | 0.113 | 0.000 | 1.000 | 1/1 |
| NUTM2A | Unknown | 0.574 | 0.001 | 0.000 | 0/3 |
| PALB2 | tumor necrosis factor; apoptosis | 0.507 | 0.000 | 0.000 | 3/10 |
| PDCD1LG2 | T-cell proliferation; immune response | 0.506 | 0.002 | 0.006 | 2/3 |
| PDE4DIP | microtubule dynamics | 0.307 | 1.000 | 0.000 | 7/6 |
| PICALM | vesicle coat protein | 0.068 | 0.000 | 0.049 | 1/4 |
| PMS2 | DNA binding protein | 0.225 | 0.000 | 0.001 | 3/5 |
| POU2AF1 | Transcriptional coactivator; immune response | 0.106 | 0.000 | 0.002 | 1/1 |
| PRF1 | apoptosis; immune response | 0.197 | 0.000 | 0.034 | 4/5 |
| PTPRB | protein phosphatase; receptor; angiogenesis | 0.156 | 0.000 | 0.002 | 7/12 |
| PTPRC | protein phosphatase;receptor; immune response | 0.361 | 0.000 | 0.000 | 36/52 |
| RABEP1 | membrane fusion; apoptosis | 0.086 | 0.000 | 0.047 | 1/6 |
| RANBP2 | G-protein modulator; transport | 0.149 | 0.001 | 1.000 | 0/5 |
| RBM15 | RNA binding protein | 0.088 | 0.000 | 0.000 | 6/9 |
| RECQL4 | DNA-dependent ATPase; chromosome segregation | 0.273 | 1.000 | 0.001 | 4/0 |
| SDHD | transfer/carrier protein | 0.394 | 0.019 | 0.009 | 1/1 |
| SET | chaperone;phosphatase inhibitor | 0.180 | 0.002 | 0.000 | 1/3 |
| SS18 | chromatin-binding protein; transcription regulation | 0.153 | 0.000 | 0.001 | 2/2 |
| STIL | Developmental protein | 0.327 | 1.000 | 0.006 | 1/1 |
| WRN | DNA helicase; DNA repair | 0.362 | 1.000 | 0.000 | 3/2 |
| ZNF384 | KRAB box transcription factor | 0.038 | 0.001 | 0.055 | 1/1 |

### 3.2 Comparison of selection estimates across functional categories

We compared global *dN, dS* and *dN/dS* values across COSMIC categories. Genes bearing only germline mutations (i.e., associated with hereditary cancer) showed significantly higher *dN/dS* estimates than genes with only somatic mutations (i.e., associated with sporadic cancer) or with both somatic and germline mutations (Figure 2A, Table 2), mainly due to a significantly increase in *dN* (Figure S2). We also observed higher *dN/dS* values for cancer genes associated with recessive mutations than for cancer genes with dominant mutations (Figure 2B, Table 2), again due to a significantly increase in dN (Figure S2). We noticed that these two mutational categories are not independent, as 33 out of 34 (97%) genes with germline mutations are associated with recessive inheritance. On the other hand, the global *dN/dS* estimates did not show significant variation among tissue types (epithelial, leukemia/lymphoma, mesenchymal and others) (Figure 2C, Table 2), cancer role (fusion genes, oncogenes and tumor suppressor genes) (Figure 2D, Table 2) or between autosomal and X chromosomes (Figure 2E, Table 2).

**Figure 2.**
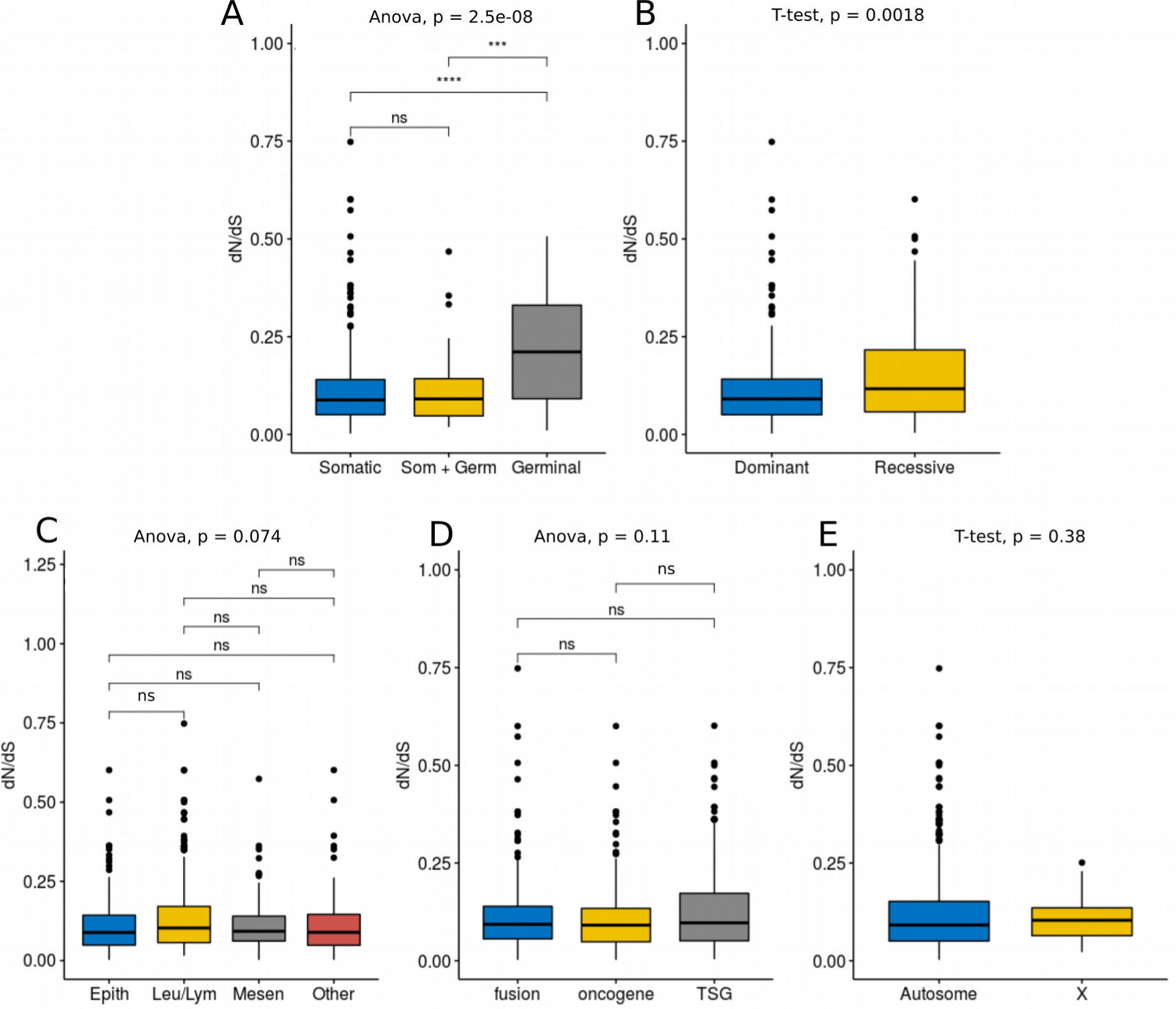
Global dN/dS across COSMIC categories. (A) mutation type, (B) genetic dominance, (C) tissue type, (D) cancer role, and (E) chromosomal type. T. ns: no significant, p-value > 0.05; (***): p-value < 0.001; (****): p-value < 0.0001.

**Table 2.**
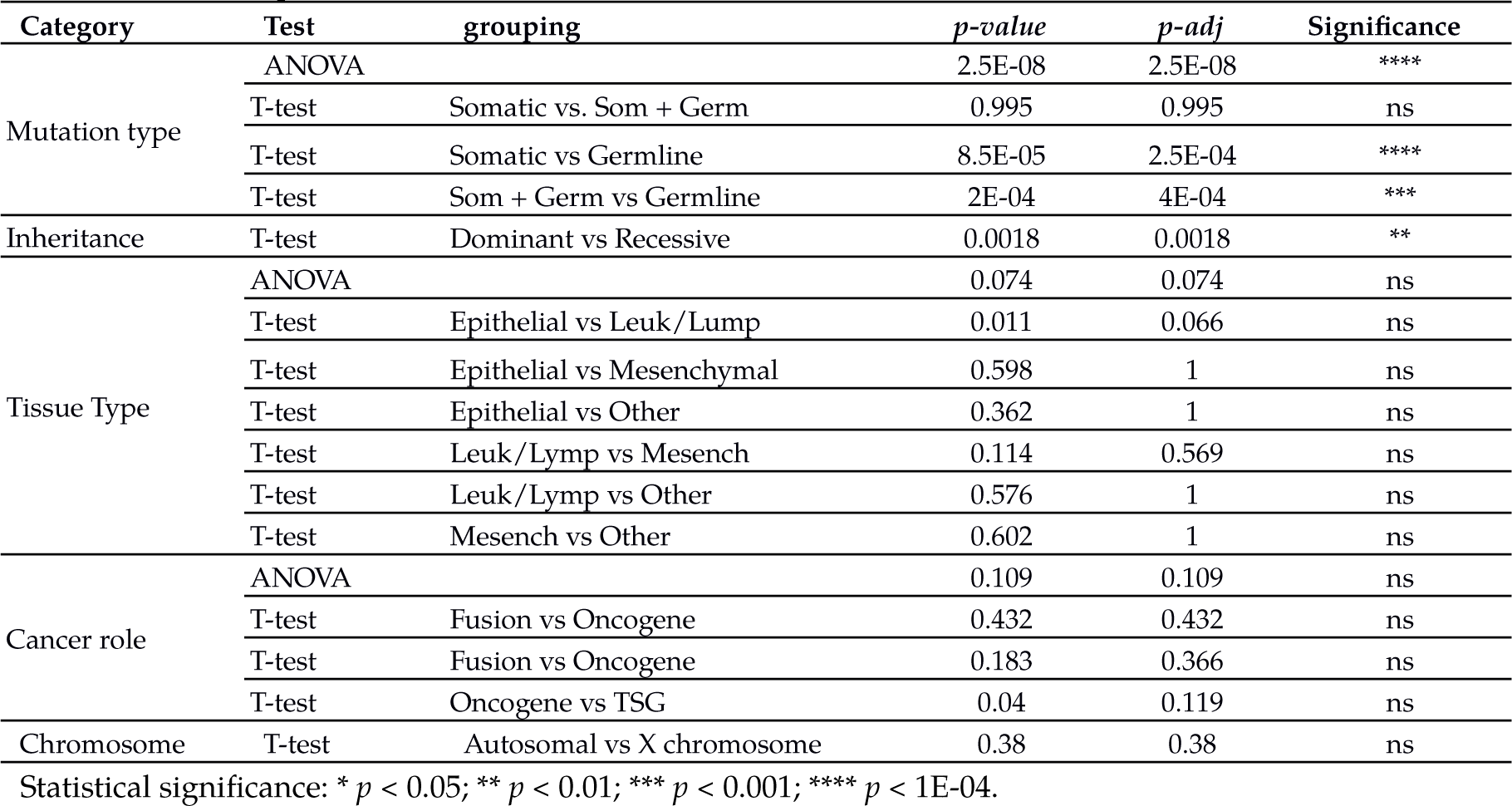
Statistical parameters of dN/dS correlations

We also compared the proportion of genes under positive selection across COSMIC categories, observing a significant increase of genes under positive selection in the germline and recessive categories (Figure 3A-B). We did not detect significant differences in the proportion of positively selected genes among tissue types, cancer role or chromosomal type (Figure 3C-E).

**Figure 3.**
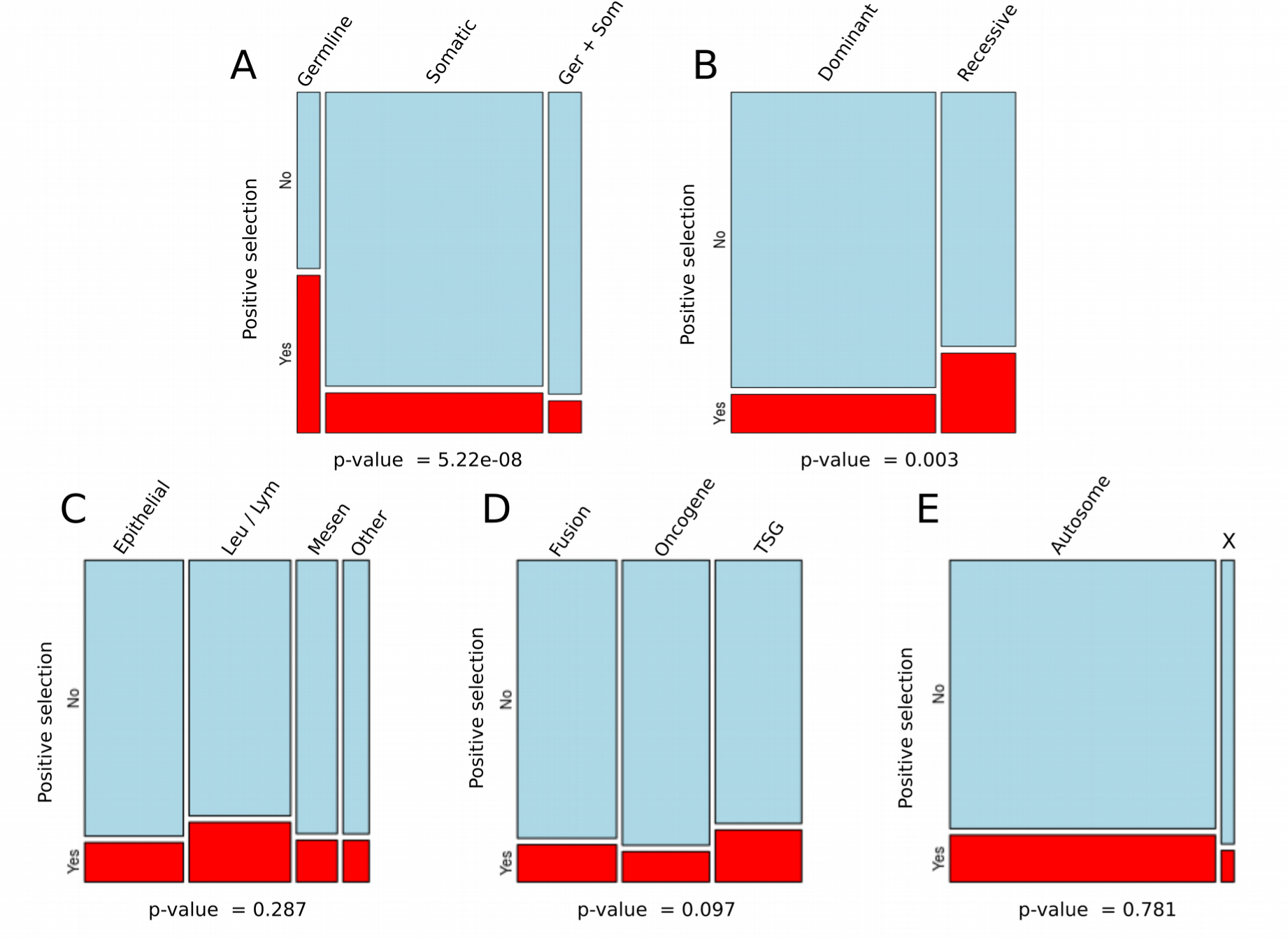
Proportion of positively selected genes across COSMIC categories. (A) mutation type, (B) genetic dominance, (C) tissue type, (D) cancer role, and (E) chromosomal type. P-values for chi-squared tests are shown underneath (significance *p* < 0.01).

### 3.3 Functional enrichment of positively selected cancer genes

The 63 cancer genes inferred to be under positive selection were enriched in biological processes associated with DNA repair, in particular interstrand crosslinking repair (Table 3). We detected a high representation in this group of members of the Fanconi Anemia Complementation Group (FANCD2, FANCG, FANCA, FANCE and FANCC), which participate with BRCA1 and BRCA2 in homologous–recombination DNA repair [33]. As mentioned above, BRCA2 yielded a strong signature of positive selection, showing the second highest number of PSSs (Table 1). We also observed an enrichment of genes that encode for proteins located in the early endosome membrane and involved in bubble DNA binding (Table 3). In addition, we observed a large portion of positively selected genes (12 out of 63, 19%, Table 1) involved in immune response, although without reaching statistical significance.

**Table 3.** Ontology terms enriched for positively selected cancer genes.

| GO class | GO term | Description | P-value | FDR | Enrichment | Genes |
| --- | --- | --- | --- | --- | --- | --- |
| Biological process | GO:0006281 | DNA repair | 1.82E-4 | 1E0 | 2.58 | BRCA2<br>BRIP1<br>CHEK2<br>DDB2<br>ERCC5<br>FANCE<br>FANCC<br>FANCD2<br>FANCG<br>PALB2<br>PMS2<br>RECQL4<br>WRN |
|  | GO:0036297 | Interstrand cross-link repair | 5.85E-4 | 1E0 | 5.71 | FANCE<br>FANCC<br>FANCD2<br>FANCG |
| Molecular function | GO:0000405 | Bubble DNA binding | 6.05E-4 | 6.61E-1 | 7.30 | ERCC5<br>RECQL4<br>BLM<br>WRN |
| Cellular component | GO:0031901 | Early endosome membrane | 5.85E-4 | 4.46E-1 | 5.71 | EGFR<br>RABEP1<br>NTRK1<br>EPS15<br>B2M |
|  | GO:0043240 | Fanconi anaemia nuclear complex | 6.05E-4 | 2.31E-1 | 7.30 | FANCE<br>FANCC<br>FANCG<br>FANCA |

### 3.4 Co-location of positively selected sites and pathogenic germline variants in BRCA2

To evaluate the potential impact of selected amino acid replacements in BRCA2, we compared the location of PSSs with that of known pathogenic germline variants [28] (Figure 4A). We detected that the stop-gained pathogenic variant Y792* in the N-terminal segment is flanked by two PSSs (Y748 and G800). We also observed several clusters of PSSs together with pathogenic variants across the BRC repeats involved in the interaction with RAD51. The residue C1159, between BRC1 and BRC2, is also in close proximity to the pathogenic variant Q1037*. In addition, we found two PSSs located in the interacting region with the polymerase Eta (POLH): the only selected residue within a BRC, E1550 in BRC2, which sits close to the truncating variant E1518*, and the residue A1708, which maps to the same segment (between BRC5 and BRC6) of the pathogenic mutation Y1762*. Furthermore, we detected a clustering of three PSSs and two pathogenic germline variants in the intervening regions between the BRC6-BRC7-BRC8 repeats. We noticed that in the site C1913, falling between BRC6 and BRC7, 11 out of 27 mammals species, including humans, have a cysteine (Cys) residue, whereas eight species carry a histidine (His). Because Cys and His residues are often involved in specific functions [34], replacements in these positions could alter protein structure and functionality. Moreover, we identified a clustering of PSSs and pathogenic variants in the domain interacting with the proteasome complex subunit SEM1 (Figure 4A-B). Interestingly, a PSS in the helical subdomain, S2572, is located in the binding interface with SEM1 (Figure 4C). Finally, nine putative PSSs mapped to a disordered segment of the C-terminal region (positions 3395-3440) with no documented activity. Among these, we observed two Cys residues whose variation could have structural implications in the protein.

**Figure 4.**
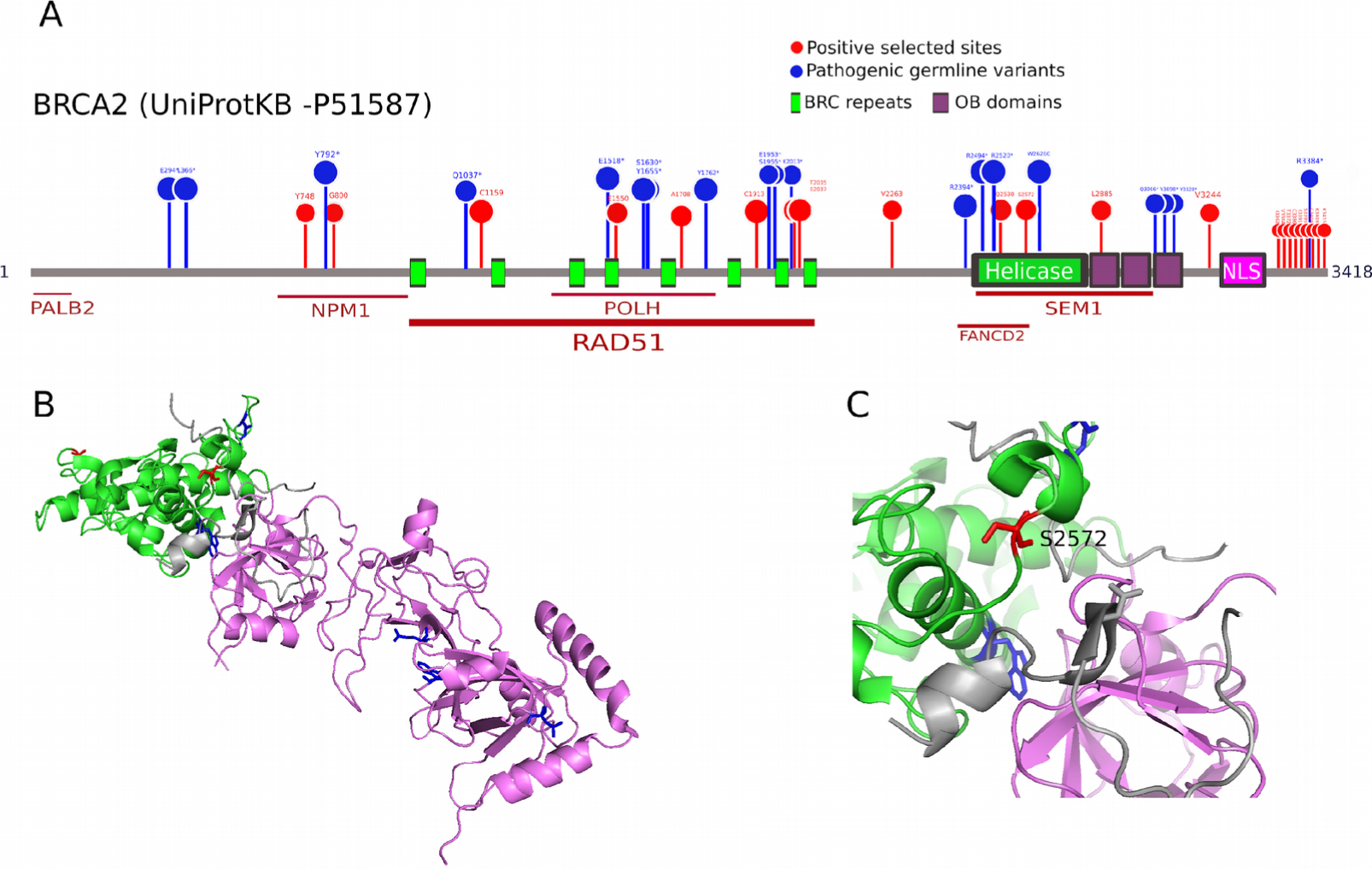
Positive selection in BRCA2. (A). BRCA2 domains with positively selected sites (PSSs; red) and pathogenic germline variants (missensense and stop-gained; blue). Proteins interacting with different domains of BRCA2 are indicated underneath. (B). Crystal structure of the rat BRCA2-SEM1 complex (PDB: 1iyj, Yang et al. 2002). Helical and OB domains are colored in green and purple, respectively, with PSS (red) and pathogenic germline variants (blue). C. BRCA2-SEM1 interface binding in the helicase domain. The S2572 residue under positive selection is labelled.

### 3.5. Strong positive selection in the extracellular region of PTPRC

We also analyzed in detail the location of PSSs within PTPRC, the gene with the highest number of such sites (Table 1). PTPRC, also known as CD45, encodes a tyrosine phosphatase that regulates T- and B-cell antigen receptor signaling and is associated with oncogenic transformation through changes in expression [35]. Remarkably, all the PSSs concentrated on the extracellular region, whereas in the cytoplasmic segment, which contains the catalytic domains, we did not detect any PSS (Figure 5A). Within the extracellular region, the PSSs clustered in the cysteine-rich (CR) and the three Fibronectin type 3 (FN3) domains, involved in cell adhesion (Figure 5B).

**Figure 5.**
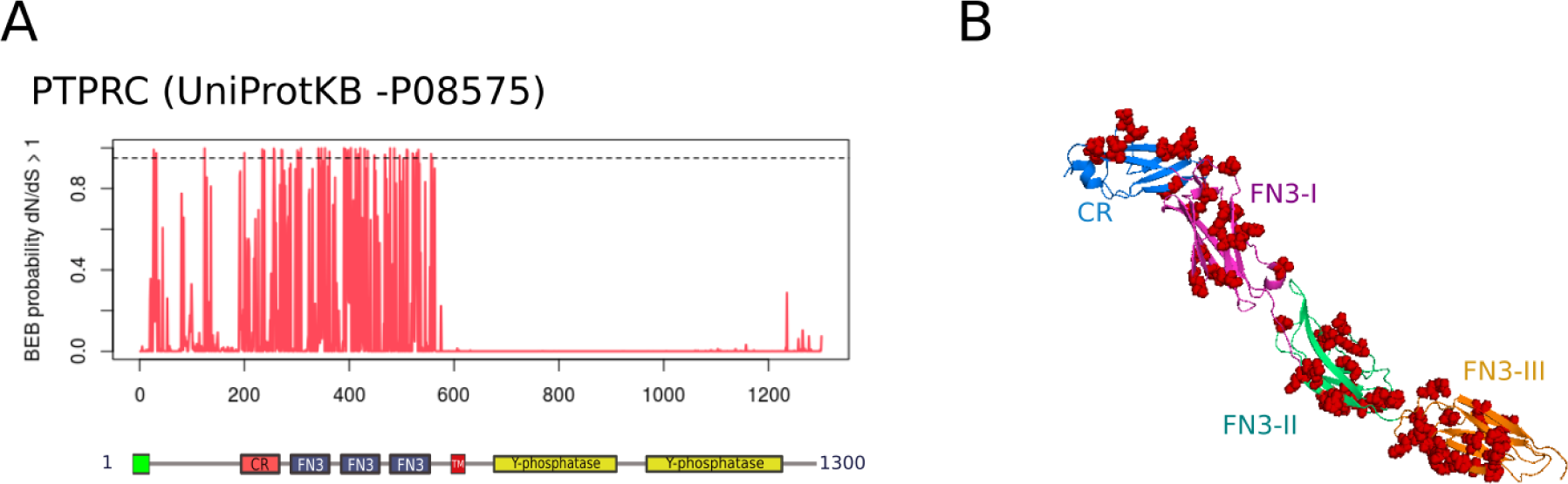
Positive selection in PTPRC. (A). BEB posterior probabilities of *dN/dS* > 1 under model M2a are represented for each residue. The dashed line indicates a BEB probability of 0.95. Protein domains are shown underneath. (B). Crystal structure of the extracellular region of PTPRC (PDB ID: 5fmv, Chang et al. *2*016), with CR and FN3 domains with different colours. PSSs are shown in red. Domain abbreviations: CR: Cysteine-rich; FN3: fibronectin 3; TM: transmembrane.

## 4. Discussion

### 4.1. Cancer genes show low dN/dS values

On average, the estimated *dN/dS* ratio across cancer genes (0.12) was somehow lower than previous estimates from mammalian genomes, which yielded values between 0.15-0.22 [7,36,37]. Our results are in concordance with Thomas et al. [32], who used pairwise comparisons between human and rodent and found a stronger purifying selection on cancer-related genes (*dN/dS* = 0.079) in comparison to other disease-related genes (*dN/dS* = 0.101) and non-disease-related genes (*dN/dS* = 0.1). Our results also concord with those from Blekhman et al. [38], who found significantly lower *dN/dS* values in genes associated with cancer between human and macaque. Because cancer genes are generally involved in essential cellular functions such as DNA repair, regulation of cell cycle and apoptosis [39], strong purifying negative selection removing deleterious germline mutations is expected.

### 4.2. Positive selection on human genes associated with hereditary cancer

Cancer genes bearing only germline mutations yielded higher *dN/dS* ratios and were more often positively selected than cancer genes with only somatic mutations. This result suggests that genes associated with hereditary cancer have less selective constraints than those genes related to sporadic cancer. Indeed, in genes under weak purifying selection it is expected that slightly deleterious variants reach higher frequencies in the populations, either by positive selection or genetic drift, than variants in genes under strong purifying selection [40]. Therefore, it is expected that genes under relaxed purifying selection are more likely associated with hereditary than with sporadic cancer. In addition, since cancer is a complex and late-onset disease, where each allele contributes to a small fraction of cancer risk and with small effects on fitness, it is plausible that genes directly associated with cancer susceptibility are under weak purifying selection [38,41,42]. In addition, genes associated with complex diseases typically are under widespread purifying selection but also show signatures of positive selection [38], in concordance with our results. On the other hand, strong purifying selection on genes causing sporadic cancer is expected, as mutations directly responsible for cancer (i.e., somatic driver mutations) should be highly deleterious.

At the same time, genes associated with recessive mutations in human cancer also showed significantly higher *dN/dS* values and were more often positively selected than genes associated with dominant mutations. However, this is result is not independent from the one just discussed, as almost all cancer genes with germline mutations in our dataset were associated with recessive inheritance. Because genes associated with germline mutations and dominant inheritance are unrepresented in our dataset, we were unable to test the direct impact of inheritance on the evolution of human cancer genes.

### 4.3 Lack of significant variation in selection across tissue, role or chromosomal type

Our analysis did not reveal significant differences in the *dN/dS* ratios or the proportion of positively selected genes among tissue types (epithelial, leukemia/lymphoma, mesenchymal and others), cancer role (fusion genes, oncogenes and tumor suppressor genes) or between autosomal and X chromosomes. Overall, this might suggest that selective pressures on human cancer genes are not directly related to cancer (see next section for further discussion). Although selection is expected to work differently in autosomal and X-linked genes because of their distinct effective population size [43], we did not identify significant differences. However, the number of X chromosome genes compared (20) was too low to guarantee strong conclusions in this regard.

### 4.4 Signalling pathways and biological functions of cancer genes under positive selection

Within the putative list of cancer genes under positive selection we found an enrichment of genes involved in the FA/BRCA pathway. Widespread positive selection on mammal genes in this pathway (specifically in BRCA2, CHEK2, FANCC, FANCB, FANCD2 and FANCE) was previously detected in mammals [44]. In addition to these genes, we also identified signatures of positive selection in FANCA and FANCG. The systematic positive selection observed on the FA/BRCA complex might suggest a mechanism of coevolution to maintain the interactions among partners of this network [44,45]. Positive selection on the FA/BRCA complex could be driven by different evolutionary mechanisms. By the one hand, positive selection on this DNA repair pathway could result in a molecular mechanism of tumor resistance to counteract the increased cancer risk associated with longevity [46]. Episodes of positive selection of some FA/BRCA components have been identified in long-lived, and cancer-resistant species. Adaptive evolution of BRCA2 has been identified in the bats *Myotis lucifugus* and *Myotis davidii* [47,48], whereas some FANC members, such as FANCA, FANCE and FANCL, were inferred to be under positive selection in the naked mole rat [47]. On the other hand, germline variants in the FA/BRCA repair pathway have been associated with hereditary breast-ovarian cancer and Fanconi Anemia (FA) in humans [33]. Therefore, it is possible that, at least a portion of selected alleles in the FA/BRCA pathway could be associated with higher cancer and FA risk. As we mentioned in the introduction, evolutionary conflicts associated with organismal fitness could be a general mechanism shaping the molecular adaptation of human cancer genes [5,8,9,11]. In the case of BRCA1 and BRCA2, these genes are involved in DNA repair also during early development, and BRCA1/BRCA2 alleles affect fetal survival in a sex-dependent manner in mammals [49]. Therefore, positive selection on BRCA1-BRCA2 genes has been previously associated with sexually antagonistic effects due to repair-proliferation tradeoff, with a pleiotropic effect on (higher) cancer hereditary risk [5,50]. Future biochemical characterization of mutants and genetic association studies will help to better understand the causes and consequences of positive selection on the components of the FA/BRCA pathway on human populations.

Immune response, placentation and spermatogenesis, expected to be shaped by pathogen-host coevolution, maternal-foetal interactions and sexual selection, respectively, are biological processes often associated with positive selection on cancer genes [5,7–10]. Interestingly, we observed a high proportion of immunes genes –albeit not statistically significant– under positive selection, which would support the contribution of pathogen-host coevolutionary interactions on driving the adaptation of human cancer genes in mammals. At the same time, the identification of several positively selected genes with prominent expression in testis (such as BRIP1, BUB1B, KTN1 and RANBP2) is concordant with the hypothesis that the pathways involved in spermatogenesis, which often evolve in response to sexual selection and intrasexual conflict [51,52], often coincide with those used by cancer cells to increase their survival and replication [7,10,53].

### 4.5 Association of residues under positive selection with pathogenic variants

We identified several genes under positive selection that have been associated with pathogenic germline variants in cancer [28] (Yuang et al. 2018). We focused on BRCA2, and observed that PSSs in this protein are localized close to pathogenic germline variants in humans. A similar distribution of sites under positive selection in BRCA2 was previously identified by O’Connell [44], who also showed an accumulation of PSSs among the BRC repeats. The residue C1159 between BRC1 and BRC2 repeats was also previously identified under positive selection in primates [54]. We identified PSSs in essential functional domains, such as SEM1 binding interface, suggesting that positive selection could affect BRCA2 function. Further examination of PSSs in conservative positions of functional domains could lead to the identification of new cancer-associated mutations.

## 5. Conclusions

In our study, genes involved in hereditary cancer showed weaker purifying selection and/or stronger positive selection than those involved in sporadic cancer. Further studies examining the interspecific variation of cancer genes in modern human populations will be essential to elucidate the contribution of long-term adaptation to human cancer inheritance. Our results suggest that, at least in mammals, positive selection acting on human cancer genes drives adaptive changes in traits related to organismal fitness, rather than select for biological functions directly related to cancer. Future studies integrating molecular data, life history traits and pathogenicity information will help to discern the selective forces behind the long-term adaptation of human cancer genes, as well as to determine the genetic conflicts between development pathways and cancer risk.

### Data availability

The data and code required to reproduce the results of this study is available in https://github.com/avicens/canger_genes_selection.

### Supplementary Materials

Figure S1: Mammal phylogenetic tree of mammalian species assembled for this study, Figure S2: Global dN and dS values according to mutation type (A, B) and inheritance (C, D), Table S1; List of Cancer Gene Census of COSMIC genes used in the study. Table S2: Genes discarded from the analysis in the different pre-processing steps, Table S3: Genes discarded from the analysis in the different pre-processing steps, Table S4: Protein Ensembl accessions of all orthologues retrieved in this study, Table S5: Evolutionary parameters estimated for each gene.

### Author Contributions

Conceptualization, A.V. and D.P.; Methodology, A.V. and D.P.; Formal Analysis, A.V.; Investigation, A.V. and D.P.; Writing-Original Draft Preparation, A.V.; Writing-Review & Editing, D.P.; Supervision, D.P.

### Funding

This research was funded by the Spanish Government (Juan de la Cierva postdoctoral fellowship IJCI-2016-29550 to A.V and research grant BFU2015-63774-P to D.P) and the European Research Council (ERC-617457-PHYLOCANCER to D.P.)

## Acknowledgments

**Conflicts of Interest:** The authors declare no conflict of interest. The founding sponsors had no role in the design of the study; in the collection, analyses, or interpretation of data; in the writing of the manuscript, and in the decision to publish the results.

## References

1. Bodmer, W.; Tomilson, I. Rare genetic variants and the risk of cancer. Curr. Opin. Genet. Dev. 2010, 20, 262–267, doi:10.1016/J.GDE.2010.04.016.

2. Martincorena, I.; Raine, K. M.; Gerstung, M.; Dawson, K. J.; Haase, K.; Van Loo, P.; Davies, H.; Stratton, M. R.; Campbell, P. J. Universal Patterns Of Selection In Cancer And Somatic Tissues. bioRxiv 2017, doi:10.1101/132324.

3. COSMIC: Catalogue of Somatic Mutations in Cancer. Available online: https://cancer.sanger.ac.uk/cosmic.

4. Bailey, M. H.; Tokheim, C.; Porta-Pardo, E.; Sengupta, S.; Bertrand, D. et al. Comprehensive Characterization of Cancer Driver Genes and Mutations. Cell 2018, 173, 371–385.e18, doi:10.1016/j.cell.2018.02.060.

5. Crespi, B. J.; Summers, K. Positive selection in the evolution of cancer. Biol. Rev. Camb. Philos. Soc. 2006, 81, 407–424, doi:10.1017/S1464793106007056.

6. Clark, A. G.; Glanowski, S.; Nielsen, R.; Thomas, P. D.; Kejariwal, A.; Todd, M. a; Tanenbaum, D. M.; Civello, D.; Lu, F.; Murphy, B.; Ferriera, S.; Wang, G.; Zheng, X.; White, T. J.; Sninsky, J. J.; Adams, M. D.; Cargill, M. Inferring nonneutral evolution from human-chimp-mouse orthologous gene trios. Science 2003, 302, 1960–3, doi:10.1126/science.1088821.

7. Nielsen, R.; Bustamante, C.; Clark, A. G.; Glanowski, S.; Sackton, T. B.; Hubisz, M. J.; Fledel-Alon, A.; Tanenbaum, D. M.; Civello, D.; White, T. J.; J Sninsky, J.; Adams, M. D.; Cargill, M. A scan for positively selected genes in the genomes of humans and chimpanzees. PLoS Biol. 2005, 3, e170, doi:10.1371/journal.pbio.0030170.

8. Kosiol, C.; Vinař, T.; da Fonseca, R. R.; Hubisz, M. J.; Bustamante, C. D.; Nielsen, R.; Siepel, A. Patterns of Positive Selection in Six Mammalian Genomes. PLoS Genet. 2008, 4, e1000144, doi:10.1371/journal.pgen.1000144.

9. da Fonseca, R. R.; Kosiol, C.; Vinař, T.; Siepel, A.; Nielsen, R. Positive selection on apoptosis related genes. FEBS Lett. 2010, 584, 469–476, doi:10.1016/j.febslet.2009.12.022.

10. Kleene, K. C. Sexual selection, genetic conflict, selfish genes, and the atypical patterns of gene expression in spermatogenic cells. Dev. Biol. 2005, 277, 16–26, doi:10.1016/J.YDBIO.2004.09.031.

11. Nielsen, R. Molecular signatures of natural selection. Annu. Rev. Genet. 2005, 39, 197–218, doi:10.1146/annurev.genet.39.073003.112420.

12. Enesmbl BioMart. Available online: http://www.ensembl.org/biomart/martview/00e8233b3e766fc8b06e4fc612687201.

13. Durinck, S.; Moreau, Y.; Kasprzyk, A.; Davis, S.; De Moor, B.; Brazma, A.; Huber, W. BioMart and Bioconductor: a powerful link between biological databases and microarray data analysis. Bioinformatics 2005, 21, 3439–3440, doi:10.1093/bioinformatics/bti525.

14. Ranwez, V.; Harispe, S.; Delsuc, F.; Douzery, E. J. P. MACSE: Multiple Alignment of Coding SEquences Accounting for Frameshifts and Stop Codons. PLoS One 2011, 6, e22594, doi:10.1371/journal.pone.0022594.

15. Capella-Gutiérrez, S.; Silla-Martínez, J. M.; Gabaldón, T. trimAl: A tool for automated alignment trimming in large-scale phylogenetic analyses. Bioinformatics 2009, 25, 1972–1973, doi:10.1093/bioinformatics/btp348.

16. Kozlov, A. RAxML-ng. Available online: https://github.com/amkozlov/raxml-ng

17. Rodríguez, F.; Oliver, J. L.; Marín, A.; Medina, J. R. The general stochastic model of nucleotide substitution. J. Theor. Biol. 1990, 142, 485–501, doi:10.1016/S0022-5193(05)80104-3.

18. Yang, Z. Maximum-likelihood estimation of phylogeny from DNA sequences when substitution rates differ over sites. Mol. Biol. Evol. 1993, 10, 1396–1401, doi:10.1093/oxfordjournals.molbev.a040082.

19. Felsenstein, J. Cofidence limits on phylogenies: an approach using the bootstrap. Evolution 1985, 39, 783–791, doi:10.1111/j.1558-5646.1985.tb00420.x.

20. Huerta-Cepas, J.; Serra, F.; Bork, P. ETE 3: Reconstruction, Analysis, and Visualization of Phylogenomic Data. Mol. Biol. Evol. 2016, 33, 1635–1638.

21. Yang, Z. PAML 4: phylogenetic analysis by maximum likelihood. Mol Biol Evol. 2007, 24, 1586–91, doi:10.1093/molbev/msm088.

22. Gharib, W. H.; Robinson-Rechavi, M. The Branch-Site Test of Positive Selection Is Surprisingly Robust but Lacks Power under Synonymous Substitution Saturation and Variation in GC. Mol. Biol. Evol. 2013, 30, 1675–1686, doi:10.1093/molbev/mst062.

23. Yang, Z.; Bielawski, J. P. Statistical methods for detecting molecular adaptation. Trends Ecol. Evol. 2000, 15, 496–503.

24. Wong, W. S. W.; Yang, Z.; Goldman, N.; Nielsen, R. Accuracy and power of statistical methods for detecting adaptive evolution in protein coding sequences and for identifying positively selected sites. Genetics 2004, 168, 1041–51, doi:10.1534/genetics.104.031153.

25. Benjamini, Y.; Hochberg, Y. Controlling the false discovery rate: a practical and powerful approach to multiple testing. J. R. Stat. Soc. B 1995, 57, 289–300.

26. Yang, Z.; Wong, W. S. W.; Nielsen, R. Bayes empirical bayes inference of amino acid sites under positive selection. Mol Biol Evol. 2005, 22, 1107–18, doi:10.1093/molbev/msi097.

27. Eden, E.; Navon, R.; Steinfeld, I.; Lipson, D.; Yakhini, Z. GOrilla: a tool for discovery and visualization of enriched GO terms in ranked gene lists. BMC Bioinformatics 2009, 10, 48, doi:10.1186/1471-2105-10-48.

28. Huang, K.; Mashl, R. J.; Wu, Y.; Ritter, D. I.; Wang, J. et al.. Pathogenic Germline Variants in 10,389 Adult Cancers. Cell 2018, 173, 355–370.e14, doi:10.1016/J.CELL.2018.03.039.

29. Kassambara, A. ggpubr: ‘ggplot2’ Based Publication Ready Plots. Available online: https://github.com/kassambara/ggpubr

30. Team, R. core R: A language and Environment for statistical computing. Available online: http://www.r-project.org.

31. Makalowski, W.; Boguski, M. S. Evolutionary parameters of the transcribed mammalian genome: An analysis of 2.820 orthologous rodent and human sequences. Proc. Natl. Acad. Sci. 1998, 95, 9407–9412.

32. Thomas, M. A.; Weston, B.; Joseph, M.; Wu, W.; Nekrutenko, A.; Tonellato, P. J. Evolutionary Dynamics of Oncogenes and Tumor Suppressor Genes: Higher Intensities of Purifying Selection than Other Genes. Mol. Biol. Evol. 2003, 20, 964–968, doi:10.1093/molbev/msg110.

33. Bogliolo, M.; Surrallés, J. The Fanconi Anemia/BRCA Pathway: FANCD2 at the Crossroad between Repair and Checkpoint Responses to DNA Damage. 2013.

34. Dayhoff, M.; Schwartz, R.; Orcutt, B. Atlas of protein sequence and structure; Dayhoff, M., Ed.; National b.; Washington DC, 1978;

35. Yuan, L.; Zeng, G.; Chen, L.; Wang, G.; Wang, X.; Cao, X.; Lu, M.; Liu, X.; Qian, G.; Xiao, Y.; Wang, X. Identification of key genes and pathways in human clear cell renal cell carcinoma (ccRCC) by coexpression analysis. Int. J. Biol. Sci. 2018, 14, 266–279, doi:10.7150/ijbs.23574.

36. Romiguier, J.; Ranwez, V.; Douzery, E. J. P.; Galtier, N. Genomic Evidence for Large, Long-Lived Ancestors to Placental Mammals. Mol. Biol. Evol. 2013, 30, 5–13, doi:10.1093/molbev/mss211.

37. Figuet, E.; Nabholz, B.; Bonneau, M.; Mas Carrio, E.; Nadachowska-Brzyska, K.; Ellegren, H.; Galtier, N. Life History Traits, Protein Evolution, and the Nearly Neutral Theory in Amniotes. Mol. Biol. Evol. 2016, 33, 1517–1527, doi:10.1093/molbev/msw033.

38. Blekhman, R.; Man, O.; Herrmann, L.; Boyko, A. R.; Indap, A.; Kosiol, C.; Bustamante, C. D.; Teshima, K. M.; Przeworski, M. Natural Selection on Genes that Underlie Human Disease Susceptibility. Curr. Biol. 2008, 18, 883–889, doi:10.1016/J.CUB.2008.04.074.

39. Futreal, P. A.; Coin, L.; Marshall, M.; Down, T.; Hubbard, T.; Wooster, R.; Rahman, N.; Stratton, M. R. A census of human cancer genes. Nat. Rev. Cancer 2004, 4, 177–183, doi:10.1038/nrc1299.

40. Eyre-Walker, A.; Keightley, P. D.; Smith, N. G. C.; Gaffney, D. Quantifying the Slightly Deleterious Mutation Model of Molecular Evolution. Mol. Biol. Evol. 2002, 19, 2142–2149, doi:10.1093/oxfordjournals.molbev.a004039.

41. Wright, A.; Charlesworth, B.; Rudan, I.; Carothers, A.; Campbell, H. A polygenic basis for late-onset disease. Trends Genet. 2003, 19, 97–106, doi:10.1016/S0168-9525(02)00033-1.

42. Spataro, N.; Rodríguez, J. A.; Navarro, A.; Bosch, E. Properties of human disease genes and the role of genes linked to Mendelian disorders in complex disease aetiology. Hum. Mol. Genet. 2017, 26, ddw405, doi:10.1093/hmg/ddw405.

43. Wolf, J. B. W.; Künstner, A.; Nam, K.; Jakobsson, M.; Ellegren, H. Nonlinear dynamics of nonsynonymous (dN) and synonymous (dS) substitution rates affects inference of selection. Genome Biol Evol. 2009, 2009, 308–19, doi:10.1093/gbe/evp030.

44. O’Connell, M. J. Selection and the Cell Cycle: Positive Darwinian Selection in a Well-Known DNA Damage Response Pathway. J. Mol. Evol. 2010, 71, 444–457, doi:10.1007/s00239-010-9399-y.

45. Qian, W.; Zhou, H.; Tang, K. Recent Coselection in Human Populations Revealed by Protein–Protein Interaction Network. Genome Biol. Evol. 2015, 7, 136–153, doi:10.1093/gbe/evu270.

46. Tollis, M.; Schiffman, J. D.; Boddy, A. M. Evolution of cancer suppression as revealed by mammalian comparative genomics. Curr. Opin. Genet. Dev. 2017, 42, 40–47, doi:10.1016/J.GDE.2016.12.004.

47. Morgan, C. C.; Mc Cartney, A. M.; Donoghue, M. T. A.; Loughran, N. B.; Spillane, C.; Teeling, E. C.; O’Connell, M. J. Molecular adaptation of telomere associated genes in mammals. BMC Evol. Biol. 2013, 13, doi:10.1186/1471-2148-13-251.

48. Zhang, G.; Cowled, C.; Shi, Z.; Huang, Z.; Bishop-Lilly, K. A.; Fang, X.; Wynne, J. W.; Xiong, Z.; Baker, M. L.; Zhao, W.; Tachedjian, M.; Zhu, Y.; Zhou, P.; Jiang, X.; Ng, J.; Yang, L.; Wu, L.; Xiao, J.; Feng, Y.; Chen, Y.; Sun, X.; Zhang, Y.; Marsh, G. A.; Crameri, G.; Broder, C. C.; Frey, K. G.; Wang, L.-F.; Wang, J. Comparative Analysis of Bat Genomes Provides Insight into the Evolution of Flight and Immunity. Science (80-.). 2013, 339, 456–460, doi:10.1126/science.1230835.

49. Healey, C. S.; Dunning, A. M.; Dawn Teare, M.; Chase, D.; Parker, L.; Burn, J.; Chang-Claude, J.; Mannermaa, A.; Kataja, V.; Huntsman, D. G.; Pharoah, P. D. P.; Luben, R. N.; Easton, D. F.; Ponder, B. A. J. A common variant in BRCA2 is associated with both breast cancer risk and prenatal viability. Nat. Genet. 2000, 26, 362–364, doi:10.1038/81691.

50. Breivik, J.; Gaudernack, G. Resolving the evolutionary paradox of genetic instability: a cost-benefit analysis of DNA repair in changing environments. FEBS Lett. 2004, 563, 7–12, doi:10.1016/S0014-5793(04)00282-0.

51. Turner, L. M.; Chuong, E. B.; Hoekstra, H. E. Comparative analysis of testis protein evolution in rodents. Genetics. 2008, 179, 2075–89, doi:10.1534/genetics.107.085902.

52. Good, J. M.; Nachman, M. W. Rates of protein evolution are positively correlated with developmental timing of expression during mouse spermatogenesis. Mol Biol Evol. 2005, 22, 1044–52, doi:10.1093/molbev/msi087.

53. Kouprina, N.; Mullokandov, M.; Rogozin, I. B.; Collins, N. K.; Solomon, G.; Otstot, J.; Risinger, J. I.; Koonin, E. V; Barrett, J. C.; Larionov, V. The SPANX gene family of cancer/testis-specific antigens: rapid evolution and amplification in African great apes and hominids. Proc. Natl. Acad. Sci. U. S. A. 2004, 101, 3077–82, doi:10.1073/pnas.0308532100.

54. Lou, D. I.; McBee, R. M.; Le, U. Q.; Stone, A. C.; Wilkerson, G. K.; Demogines, A. M.; Sawyer, S. L. Rapid evolution of BRCA1 and BRCA2 in humans and other primates. BMC Evol. Biol. 2014, 14, 155, doi:10.1186/1471-2148-14-155.

